# An interplay of population size and environmental heterogeneity explains why fitness costs are rare

**DOI:** 10.1101/2020.10.26.355297

**Authors:** Yashraj Chavhan, Sarthak Malusare, Sutirth Dey

**Affiliations:** Indian Institute of Science Education and Research (IISER) Pune, Dr Homi Bhabha Road, Pashan, Pune, Maharashtra, 411008, India; Gaia Doctoral School, Institut des sciences de l’évolution (ISEM) 1093-1317 Route de Mende, 34090, Montpellier, France

**Keywords:** Costs of adaptation, whole-genome whole-population sequencing, mutation supply, fluctuating environments, ecological specialization, experimental evolution, maladaptation, mutation fixation

## Abstract

Theoretical models of ecological specialization commonly assume that adaptation to one environment leads to fitness reductions (costs) in others. However, empirical studies often fail to detect such costs. We addressed this conundrum using experimental evolution with *Escherichia coli* in several homogeneous and heterogeneous environments at multiple population sizes. We found that in heterogeneous environments, smaller populations paid significant costs, but larger ones avoided them altogether. Contrastingly, in homogeneous environments, larger populations paid more costs than the smaller ones. Overall, large population sizes and heterogeneous environments led to cost avoidance when present together but not on their own. Whole-genome whole-population sequencing revealed that the enrichment of multiple mutations within the same lineage (and not subdivision into multiple distinct specialist subpopulations) was the mechanism of cost avoidance. Since the conditions revealed by our study for avoiding costs are widespread, it explains why the costs expected in theory are rarely detected in experiments.

## Introduction

Costs of adaptation, also known as ‘fitness costs’ and ‘true trade-offs,’ entail that a fitness increase in one environment leads to a fitness decline in another (Bono *et al*. 2017). Such costs are instrumental in understanding why species tend to favour a particular set of environmental conditions over others (Fry 1996; Bono *et al*. 2017). Apart from answering such fundamental questions in evolutionary ecology, understanding fitness costs can also help in combating practical challenges like the rampant spread of antibiotic resistance(Andersson & Hughes 2010) and forecasting how species would respond to climate change (Wallenstein & Hall 2012). Although such costs are a fundamental assumption of numerous models of ecological specialization (Levins 1968; Futuyma & Moreno 1988; Fry 1996), a large number of experimental evolution studies spanning diverse taxa have failed to detect them (Rausher 1984; Coustau *et al*. 2000; Vasilakis *et al*. 2009; Vila-Aiub *et al*. 2009; Friman & Buckling 2013). Consequently, explaining this rarity of detectable fitness costs has been a major challenge for evolutionary studies over the last two decades (Joshi & Thompson 1995; Fry 1996; Agrawal *et al*. 2010; Remold 2012).

Here we investigate the evolutionary emergence and avoidance of fitness costs in asexual microbial populations, which have proven to be convenient model systems for experimental evolution studies over hundreds of generations (Kassen 2014; Bono *et al*. 2017). Whereas numerous microbial experimental evolution studies have reported the absence of detectable fitness costs altogether, several others have found such costs in some microbial populations but not in others (see Table S1 for a detailed list).

An important but trivial explanation for the failure to find fitness costs is the absence of any real costs altogether (Coustau *et al*. 2000). Indeed, some recent investigations have found the pleiotropy of new mutations to be largely positive and not negative (*i.e*., costly) (Sane *et al*. 2018). More importantly, the extant literature offers three distinct explanations as to why fitness costs may exist but remain undetected in empirical studies (Velicer & Lenski 1999; Coustau *et al*. 2000). First, costs can be detected only under certain environmental conditions which the experimental setup may fail to provide (Coustau *et al*. 2000; Agrawal *et al*. 2010; Kassen 2014). Second, it is a statistically demanding task to detect negative pleiotropy (aka antagonistic pleiotropy), the very foundation of fitness costs, which entails that a mutation that is beneficial in one environment is deleterious in another. This is because the statistical significances of both the beneficial and deleterious effects need to be established simultaneously for detecting costs. If the experiment does not have enough statistical power to detect these opposite effects simultaneously, costs would not be detected (Coustau *et al*. 2000; Bono *et al*. 2017). Third, the emergence of fitness costs is expected to require a threshold amount of time; such costs may appear only after several thousand generations of microbial evolution have passed (Velicer & Lenski 1999; Jasmin & Zeyl 2013; Satterwhite & Cooper 2015), and therefore would be detectable only in very long-term experimental evolution studies.

A recent meta-analysis of microbial experimental evolution studies provides a new explanation for the emergence of fitness costs based on environmental heterogeneity, suggesting that environments imposing a single (homogeneous) selection pressure frequently lead to fitness costs that can be avoided in heterogeneous environments (which fluctuate across multiple individual selection pressures) (Bono *et al*. 2017). Antagonistic pleiotropy can evolve freely if the environment does not allow the ensuing costs of adaptation to be expressed. Since selection would be blind to the antagonistic pleiotropic effects if the environment does not change, fitness costs are more likely to appear in homogeneous environments with a single selection pressure as compared to heterogeneous environments with multiple fluctuating selection pressures.

Unfortunately, the above prediction holds only weakly as many microbial experimental evolution studies have failed to find lower costs in heterogeneous environments as compared to homogeneous ones (Jasmin & Kassen 2007b; Presloid *et al*. 2008; Friman & Buckling 2013; Ketola & Saarinen 2015). This opens up the possibility that factors other than environmental heterogeneity may be important in shaping the emergence of fitness costs. One such factor is populations size, which has been shown to be important in shaping the correlated changes in populations’ fitness in alternative environments (Chavhan *et al*. 2019a, 2020). For example, a recent study showed that larger populations evolving in a homogeneous environment containing a single carbon source suffer greater fitness costs in alternative environments (Chavhan *et al*. 2020). These results could be explained with a combination of two notions. First, adaptation in very large populations is primarily driven by beneficial mutations of large effect sizes (Desai & Fisher 2007a; Chavhan *et al*. 2019b). Second, larger beneficial mutations are expected to carry heavier disadvantages in alternative environments (Lande 1983; Orr & Coyne 1992).

Taken together, the extant literature suggests that environmental heterogeneity and population size are two important factors that can potentially shape the evolution of fitness costs. However, the effects of the interaction of these two factors remains unknown. Interestingly, this interaction can be expected to play out in two contrasting ways.

First, if mutational pleiotropy across environmental components is largely antagonistic, and large benefits in one context entail large costs in another, the multiplicity of selection pressures in a heterogeneous environment would prevent the enrichment of costly large effect mutations, even if the latter were accessible to the population. This is akin to Fisher’s formulation of micromutationism where adaptation is expected to proceed via mutations of small effects(Fisher 1930). In this scenario, in heterogenous environments, both large and small populations are expected to pay similar costs.

Second, in a heterogeneous environment, evolving in larger numbers can make populations stumble upon greater number of mutations that are beneficial in a given environmental state, but not necessarily in others. The presence of multiple mutations within an individual belonging to a large asexual population has the potential to offset the costs carried by individual mutations in isolation. In this scenario, adapting in larger numbers in a heterogeneous environment would lead to the avoidance of fitness costs. Interestingly, bacterial experimental evolution studies conducted in heterogeneous environments agree with this notion: studies on smaller populations tend to detect costs, while those using larger populations do not (Fig 1a).

**Fig. 1.**
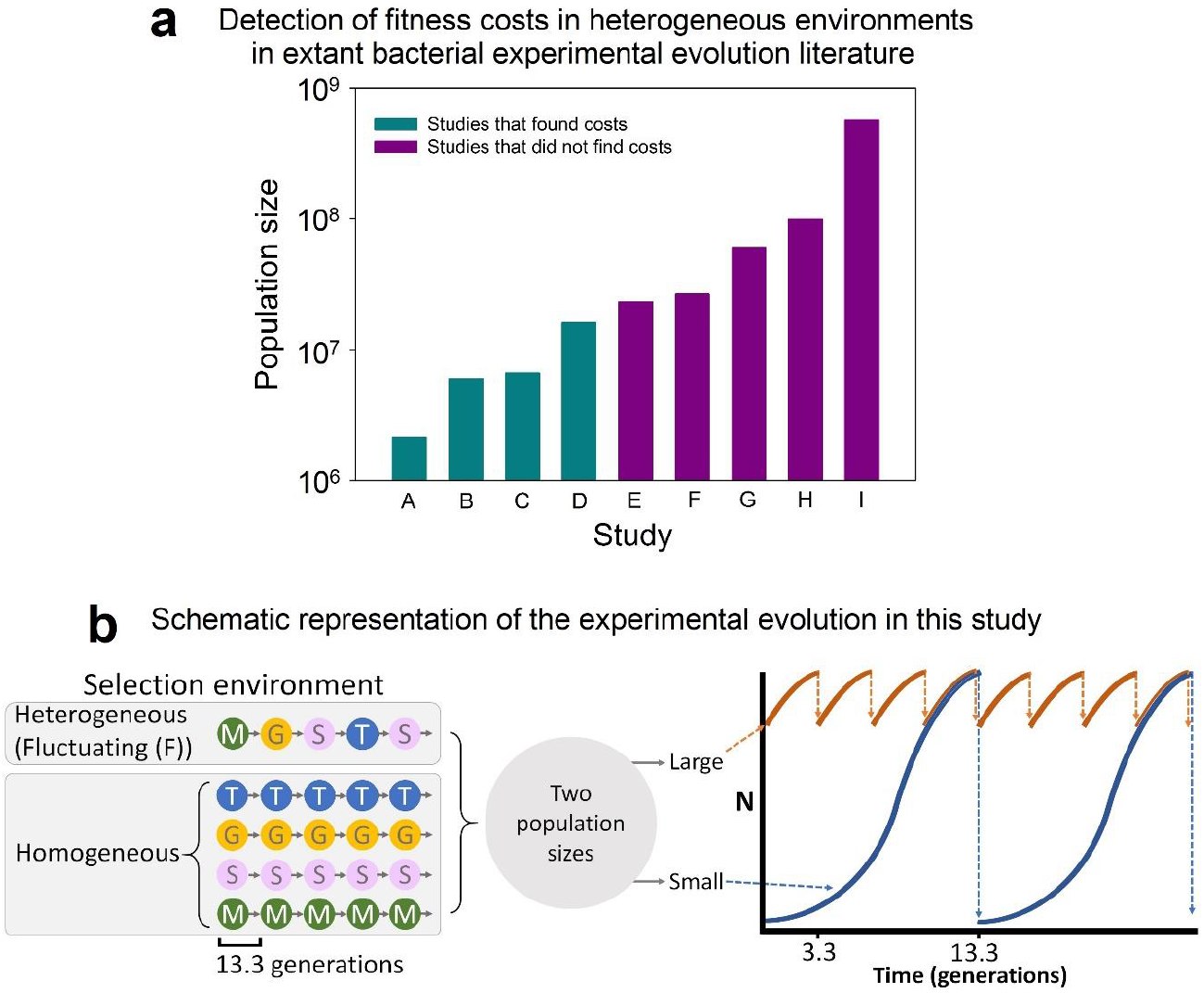
**(a)** The harmonic mean sizes of laboratory populations in existing bacterial experimental evolution studies on fitness costs conducted in heterogeneous environments. See Supplementary Text (ST.1) for the details of the studies shown in the ordinate. **(b)** A schematic representation of our evolution experiment. The experimental populations were maintained in five distinct environments at two different population sizes. T, G, S, and M refer to thymidine, galactose, sorbitol, and maltose, respectively. N stands for absolute population size. In the heterogeneous (randomly fluctuating environment, the identity of the sole carbon source changed every 13.3 generations. See the text for further details.

Stated differently, in homogeneous environments, larger populations are expected to pay heavier costs of adaptation (Chavhan *et al*. 2020). However, in heterogeneous environments, larger populations may either pay similar or lower costs as compared to smaller populations, depending upon which one of the above two possibilities dominates the underlying adaptive dynamics. To the best of our knowledge, no studies in the existing literature have tested these contrasting expectations empirically.

Here we study how environmental heterogeneity and population size interact with each other to influence the evolutionary emergence or avoidance of fitness costs. To this end, we use experimental evolution with clonally derived *Escherichia coli* populations in both heterogeneous and homogeneous environments at different population sizes for ~480 generations. We investigate if population size has similar effects on fitness costs in homogeneous and heterogeneous environments. We also test if evolving in a heterogeneous environment can lead to cost avoidance, regardless of the population size. We show that population size influences costs in opposite ways in homogeneous and heterogeneous environments. Interestingly, large population size and heterogeneous environments lead to evolutionary avoidance of costs when present simultaneously but not in isolation. Mutational frequency distributions obtained by whole-genome whole-population sequencing revealed how environmental heterogeneity led to cost avoidance in large populations but not in smaller ones. Based on these observations, we propose a new explanation for the rarity of fitness costs in evolutionary and ecological studies, which can account for several contrasting observations made in the last two decades of microbial experimental evolution.

## Results and Discussion

### Large population size and heterogeneous environments led to cost avoidance when present together but not on their own

We carried out experimental evolution with clonally derived *E. coli* populations in five different nutrient limited environmental conditions at two different population sizes for ~480 generations (Fig. 1b). This design gave rise to ten different evolutionary regimens (FL, FS, TL, TS, GL, GS, SL, SS, ML, and MS; where the first letter represents the sole carbon source in a regimen’s selection environment (fluctuating (F), thymidine (T), galactose (G), sorbitol (S), and maltose (M)) while the second letter stands for its population size (L (large) and S (small)). The harmonic mean population size for our principal treatment (F, Fluctuating (heterogeneous) environment) was ~1.01 × 10^8^ for the large (FL) populations and ~4.04 × 10^5^ for the small (FS) populations. Moreover, the adaptively relevant population sizes for L and S in this treatment were approximately equal to 9.13 × 10^6^ and 2.28 × 10^3^, respectively (Chavhan *et al*. 2019b). In the FL and FS regimens, the identity of the sole carbon source fluctuated randomly across four distinct states (T, G, S, and M) approximately every ~13.3 generations (Fig. 1b). Our study also involved four distinct homogeneous environmental controls, each with an unchanging identity of the sole carbon source corresponding to one of T, G, M, or S (Fig. 1b). With six replicates per regimen, our experiment involved 60 independently evolving populations in total. All the large (L) populations faced a periodic bottleneck ratio of 1:10 while all the small (S) populations experienced a periodic bottleneck of 1:10^4^. We manipulated the timing and frequency of bottlenecks to ensure that large and small populations did not spend significantly different times in the stationary phase (Fig. 1b; see Methods for details).

We conducted growth measurements to obtain high-resolution growth curves for all the 60 independently evolving populations in all four distinct sole carbon sources (T, G, M, and S) at the end of the evolution experiment. We used the maximum growth rate (R) as the measure of fitness (Leiby & Marx 2014; Karve *et al*. 2015; Chavhan *et al*. 2019a, b) (see Methods for details). We identified the occurrence of significant costs of adaptation in our experimental populations as cases that showed adaptation to one environment and simultaneous maladaptation to another. To this end, we carried out single sample *t*-tests with the ancestral fitness level (scaled to 1) as the reference value. We then corrected for family-wise error rates using the Holm-Šidàk procedure (Abdi 2010). Cases with fitness > 1 (corrected *P* < 0.05) were identified as adaptations; analogously, cases with fitness < 1 (corrected *P* < 0.05) were identified as maladaptations.

We found that twenty-one out of the forty possible combinations of regimen and assay environment showed significant fitness changes as compared to the common ancestor (corrected *P* < 0.05; see Table S2). We used this information to analyse the effects of two factors that are expected to be important in shaping the evolution of fitness costs in bacterial populations, namely population size and environmental heterogeneity.

In the heterogeneous (F) environment, the large populations (FL) completely avoided costs across all the environmental pairs under consideration (Fig. 2; Tables S2 and S3). FL adapted simultaneously to both T and G and did not show a significant change in fitness (vis-à-vis the common ancestor) in S and M (Fig. 2; Tables S2 and S3). On the other hand, the small populations evolved in the heterogeneous environment (FS) adapted only to T, becoming maladapted to (and hence paid a cost of adaptation in) the other three sole carbon sources (G, S, and M) (Fig. 2; Tables S2 and S3). Taken together, when evolved in the heterogeneous (F) environment, the small populations paid greater costs than the large populations, with the latter avoiding all costs altogether.

**Fig. 2.**
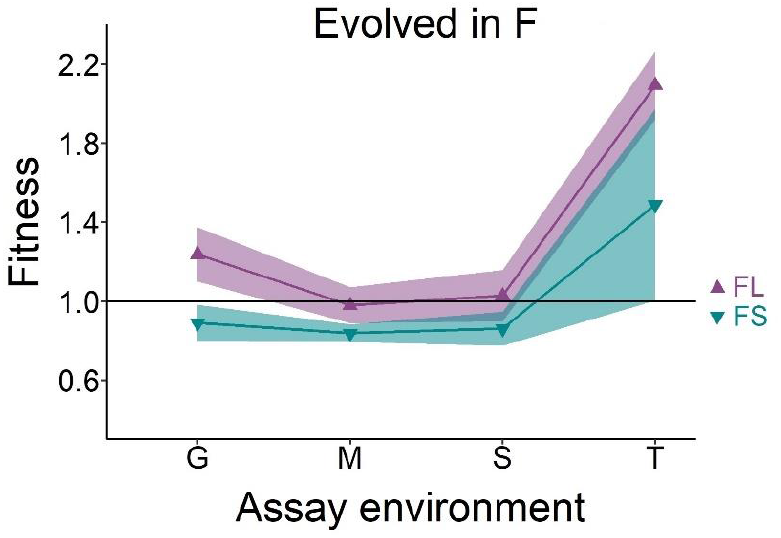
Reaction norms of fitness of large (FL) and small (FS) populations evolved in the heterogeneous environment across the four environmental states faced during evolution. G, M, S, and T represent galactose, maltose, sorbitol, and thymidine, respectively. The error bands represent 95% CI (*t*-distribution). The solid black line represents the ancestral fitness. FL adapted simultaneously to two environments (G and T) and avoided the costs of adaptation across all the environmental pairs under consideration. Contrastingly, FS adapted to T and paid costs of adaptation in the other three environments (G, M, and S). See Tables S2 and S3 for detailed statistics.

Interestingly, in the homogeneous (control) environments, the above pattern of costs reversed completely. Here, the large populations paid heavier costs of adaptation than the smaller ones (Fig. 3; the fitness changes pertaining to selection in homogeneous T and G environments have been reported previously (Chavhan *et al*. 2020)). Specifically, when evolved in homogeneous T, both TL and TS paid significant costs. Interestingly, the costs suffered by TL were significantly greater than those suffered by TS, regardless of the environmental pair in question (Fig. 3; Tables S2 and S4). When evolved in homogeneous G, only GL paid costs of adaptation (GS failed to adapt significantly to the homogeneous G selection environment). None of the populations evolved in homogeneous M and S environments adapted to their respective selection environments, regardless of the population size; hence, there were no cots of adaptation in these regimens (Fig. 3; Tables S2 and S4).

**Fig. 3.**
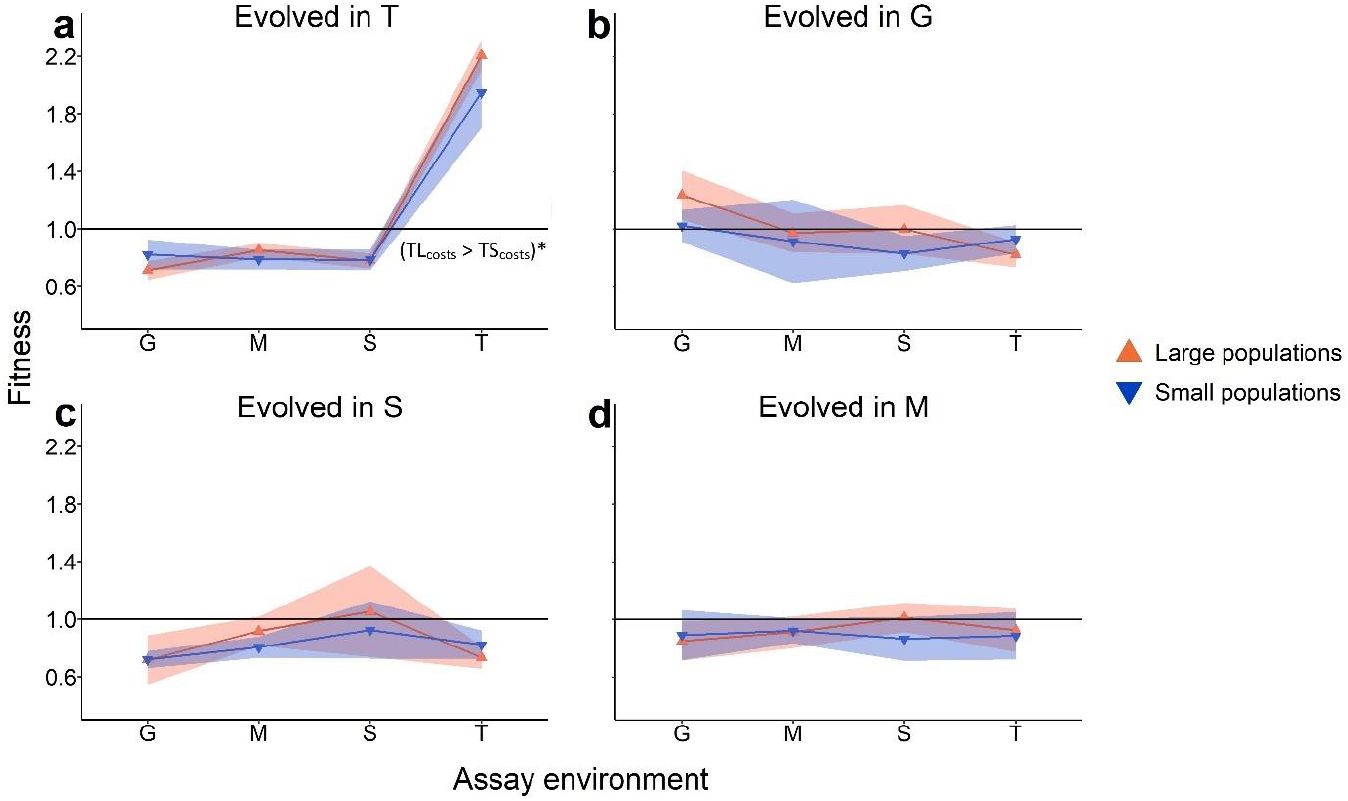
Reaction norms of fitness of populations evolved in homogeneous environments. G, M, S, and T represent galactose, maltose, sorbitol, and thymidine, respectively. The error bands represent 95% CI (*t-*distribution). The solid black line represents the ancestral level of fitness. See Tables S2 and S3 for detailed statistics. **(a)** When evolved in T, both the large (TL) and small populations paid costs in the other three environments (G, M, and S). The costs paid by TL were significantly greater than those paid by TS (Chavhan *et al*. 2020). **(b)** GL paid significant costs in T. GL did not have significantly different fitness relative to the common ancestor in M and S. GS did not adapt significantly to G. Hence there were no costs of adaptation in this case. **(c)** Both SL and SS failed to show significantly different fitness with respect to the common ancestor. Hence there were no costs of adaptation in either SL or SS. **(d)** Neither ML nor MS had significantly different fitness with respect to the common ancestor. Hence there were no costs of adaptation in either ML or MS.

Homogeneous T and G environments are known to exhibit reciprocal fitness trade-offs with each other (Chavhan *et al*. 2020). In other words, adaptation to T is accompanied by maladaptation to G, and vice-versa (Chavhan *et al*. 2020). Agreeing with this notion, we found that when evolved in the heterogeneous environment (where the sole carbon source fluctuated randomly), the small populations (FS) indeed suffered from the T-G costs. Specifically, FS adapted to T but became significantly maladapted to G (Fig. 2; Table S3). Contrastingly, the large populations evolved in the heterogeneous environment (FL) completely bypassed the expected T-G trade-off, adapting simultaneously to both the carbon sources, thereby avoiding the costs of adaptation across this environmental pair (Fig. 2, Table S3).

Taken together, evolution in the ten regimens of our study reveals that an interplay of environmental heterogeneity and population size shaped how fitness costs evolved. We found that population size had opposite effects on costs of adaptation during evolution in heterogeneous versus homogeneous environments. While in homogeneous environments, larger populations evolved greater costs; contrastingly, in heterogeneous environments, smaller populations paid greater costs while larger ones avoided them altogether. Importantly, neither environmental heterogeneity nor population size could sufficiently explain the emergence (or avoidance) of costs on their own (compare Figs. 2 and 3). Overall, costs could be avoided altogether only when heterogeneous environments and large population size were present simultaneously (the FL regimen).

### Conventional explanations cannot account for the avoidance of fitness costs in our experiments

Conventional notions about the rarity of detectable fitness costs failed to explain our observations. One such explanation is that perhaps the experiment did not provide the relevant conditions for costs to be expressed (Coustau *et al*. 2000; Agrawal *et al*. 2010; Kassen 2014). This was not the case in our experiments as several environmental pairs showed significant costs of adaptation. Another potential explanation is that the substantial statistical demands of establishing antagonistic pleiotropy were not met (Coustau *et al*. 2000; Anderson *et al*. 2013; Ågren *et al*. 2013; Bono *et al*. 2017). However, we were able to statistically detect costs caused by antagonistic pleiotropy in multiple regimens and in both homogeneous and heterogeneous environments (Figs 2 and 3). Finally, an often-quoted explanation for the lack of fitness costs is the relatively short duration of the experimental evolution study (Velicer & Lenski 1999; Jasmin & Kassen 2007a; Jasmin & Zeyl 2013; Satterwhite & Cooper 2015; Schick *et al*. 2015). However, this was simply not true in our case, as several fitness costs had already emerged over the ~480 generations of selection.

As discussed earlier, evolution in heterogeneous environments is expected to lead to lower costs than evolution in homogeneous environments because the former offer multiple dynamic selection pressures (Bono *et al*. 2017). Although our temporally heterogeneous (F) environment contained only a single carbon source at any given point of time, the identity of this carbon source fluctuated randomly over four states every ~13.3 generations. Therefore, selection was not expected to be blind to the pleiotropic fitness effects of mutations across T, G, M, and S. Despite evolving in such a heterogeneous environment, the FS populations paid significant fitness costs. Thus, Fig. 2 shows that contrary to the expectations of the extant literature (Bono *et al*. 2017), the presence of multiple dynamic selection pressures can be insufficient for cost avoidance.

Interestingly, evolutionary success in fluctuating environments is reflected by the geometric mean (GM) fitness across the states about which the environment oscillates (and not necessarily the arithmetic mean fitness) (Orr 2007; Kassen 2014). We found that across G, M, S and T, FL had significantly greater GM fitness than both FS and the common ancestor (Fig. S1a; Table S5). In contrast, the GM fitness of FS was not significantly different from the ancestral value (Fig. S1a; S5). Furthermore, as expected, evolution in homogeneous (unchanging) environments did not result in increased GM fitness above the ancestral value, regardless of the population size (Fig. S1b and Table S5). FL was better prepared to face the fluctuating environment than all the eight homogeneous environmental regimens (Tables S6 and S8) Surprisingly, the preparedness of FS to face the environmental fluctuations across G, M, S and T was similar to most homogeneous environment regimens (Tables S7 and S8). These observations highlight the key role played by population size in shaping fitness relationships across the component states of heterogeneous environments. Thus, the mere presence of multiple dynamic selective pressures in a heterogeneous environment was not enough to prevent costs of adaptation, which ultimately precluded any significant increase in the geometric mean fitness of FS.

### The genetic basis of cost avoidance

The observation that FS suffered substantial costs that were completely avoided by FL can be explained by the notion that in the presence of multiple selection pressures, a threshold amount of mutational supply is required to avoid costs. Owing to their relatively larger size, FL are expected to have much higher mutational supply as compared to FS. We hypothesised that FL enriched a larger number of mutations than FS, which made them adapt to multiple carbon sources, thereby avoiding the costs that were paid by the FS populations. To validate this hypothesis, we performed end-point whole-genome whole-population sequencing in three randomly chosen populations each from FL and FS. For our analysis, we considered only mutations that had a frequency ≥ 10% (Lang *et al*. 2013; Bailey *et al*. 2015; Copin *et al*. 2016; McDonald *et al*. 2016; Swings *et al*. 2017). Theory suggests that any mutation rising to frequencies ≥ 10% within 480 generations in any of our treatment populations is likely to be beneficial and highly unlikely to be neutral or deleterious (Desai & Fisher 2007a; Good *et al*. 2012; Cooper 2018). Consistent with this notion, we found that the number of mutations rising to frequencies ≥ 10% was much greater in FL as compared to FS (Fig. 4).

**Fig. 4.**
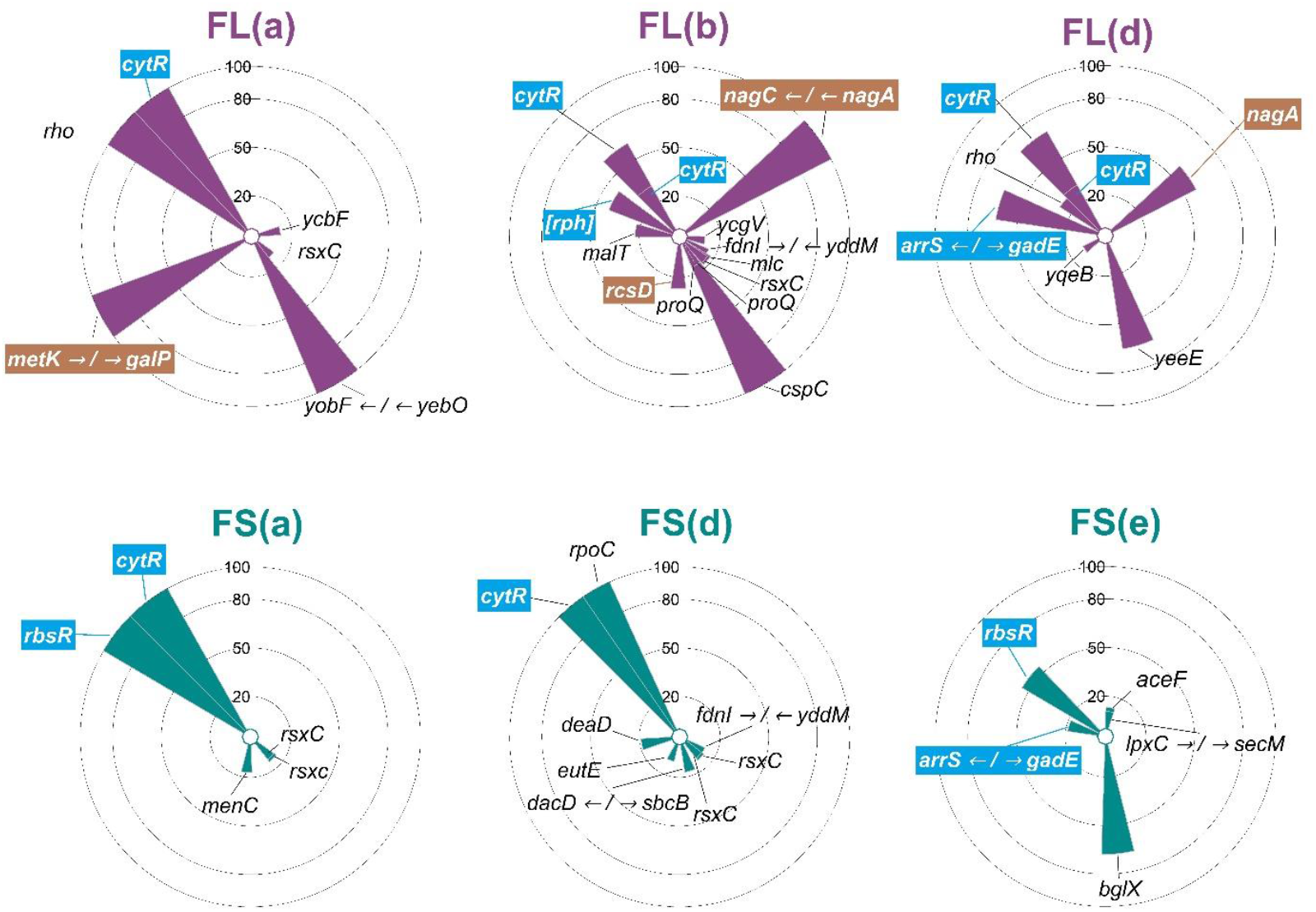
The spectrum of mutations observed in FL and FS after 480 generations. Three randomly chosen replicate populations each of FL (upper row) and FS (lower row) were subjected to whole-genome whole-population sequencing. The radial bars are located at the genomic position of the observed mutations and their heights represent the corresponding mutational frequency. The mutated loci known to be associated with thymidine (T) utilization are highlighted in blue while those associated with galactose (G) utilization are highlighted in brown. See Table S9 for details.

A detailed description of the various observed mutations is given in Table S9, while the key observations and their interpretations are described below.

We found that the loci mutated in FL are known to be associated with the uptake and/or metabolism of either G or T in the extant literature. Contrastingly, the loci mutated in FS had known links to T uptake and/or metabolism but none with that of G (Fig. 4). This agrees with the observation that FL adapted to both T and G while FS adapted to T but not to G.

Interestingly, some mutated loci that could be linked with T adaptation were common across FL and FS while others were found exclusively in either FL or FS. Remarkably, five out of the six sequenced populations (3/3 in FL and 2/3 in FS) had high frequencies of mutations in *cytR* (Fig. 4), an important regulator of thymidine metabolism that is instrumental in the regulation of pyrimidine uptake and degradation (Hammer-Jespersen & Munch-Petersen 1975; Valentin-Hansen *et al*. 1996). Similarly, insertions in the upstream regulator region of *gadE*, a transcriptional activator that plays an important role in thymidine metabolism (Ketcham 2019), were found to be enriched in one replicate of both FL and FS.

A deletion in the expressed but non-active exoribonuclease *rph* got enriched in an FL population, but was not found in any of the FS populations (Fig. 4). Such a deletions is likely to affect the expression of *pyrE*, a key gene in thymidine biosynthesis whose promoter lies within *rph* (Gama-Castro *et al*. 2016; Wytock *et al*. 2018). On the other hand, two out of the three sequenced FS populations showed mutations in *rbsR*, the ribose operon repressor that has known links to thymidine metabolism (Shimada *et al*. 2013). Interestingly, mutations in *rbsR* did not get enriched in any of the three FL populations.

The mutations identified to be associated with adaptation in Gal were found exclusively in FL (none in FS) and were distributed across diverse loci. For example, mutations influencing the expression of *nagA* and *nagC* genes were found at frequencies > 50% in two out of the three sequenced FL populations (Fig. 4). Mutations in these genes are known to increase fitness in galactose minimal media (Soupene *et al*. 2003; El Qaidi *et al*. 2009). Similarly, a mutation in the operator of *galP*, the galactose:H^+^ symporter (a gene that is instrumental in galactose uptake) got fixed in one FL population (Fig. 4).

We also found a high frequency mutation directly associated with maltose utilization in one of the FL populations, but none in FS (Fig. 4, Table S9). The presence of this mutation could explain the avoidance of costs in M that could have arisen due to T-associated mutations in FL. Furthermore, we also found several mutations in genes with widespread effects that were not specific to the uptake or metabolism of the carbon sources used in this study (Fig. 4, Table S9). Since fitness in T has been shown to be negatively correlated with fitness in G (Chavhan *et al*. 2020), mutations beneficial in T are likely to be deleterious in G, and vice versa. Hence, the presence of several T-associated mutations at high frequencies in FS can explain their maladaptation to G. Moreover, we did not find any known G-associated mutations in FS that could alleviate the putative maladaptive effects of T-associated mutations in G. Had there been no G-associated mutations in FL, the enrichment of a relatively larger number of T-associated mutations should have led to greater maladaptation of FL in G. However, we found several G-associated mutations at high frequencies in FL that can explain these populations’ adaptation to G.

The presence of both T- and G-associated mutations in FL agrees with the observation that this regimen adapted to both T and G. The large population size of FL could have allowed them to stumble upon highly rare mutations that were simultaneously beneficial in multiple environments (Li *et al*. 2019) (T and G in this case). However, the convergent enrichment of multiple mutations at the level of loci (e.g., within *cytR* and upstream of *gadE* (Fig. 4)) in FL and FS makes such a possibility unlikely. Although the investigation of such individual and epistatic effects of mutations on fitness across different environments is interesting in its own right, it is outside the scope of our study, which is primarily targeted towards unravelling the interactive effects of population size and environmental heterogeneity in shaping fitness costs.

Taken together, the genomic changes enriched during evolution in the heterogeneous environment were congruent with the phenotypic observation that the large (FL) avoided all the fitness costs that were suffered by the small (FS) populations. Having discussed the match between our phenotypic and genotypic observations, we now turn to the population genetic drivers that could have shaped evolution in our experiments.

Antagonistic pleiotropy can readily explain the positive relationship between population size and fitness costs observed in homogeneous environments (Rose & Charlesworth 1980; Cohan *et al*. 1994; Holt 1996; Cooper & Lenski 2000; Cooper 2014) (Fig. 3). Since these populations faced only one carbon source throughout the experiment, their evolution was blind to fitness changes in other carbon sources. The pleiotropic disadvantages of beneficial mutations are generally expected to be correlated with their direct effects (Lande 1983; Orr & Coyne 1992; Otto 2004; Chavhan *et al*. 2019a). Since the larger asexual populations adapt primarily via beneficial mutations with relatively greater direct effect sizes (Desai *et al*. 2007; Desai & Fisher 2007b; Sniegowski & Gerrish 2010; Chavhan *et al*. 2019b), adapting to homogeneous environments in larger numbers should lead to heavier costs of adaptation, as observed in our study (Fig. 3) (Chavhan *et al*. 2020).

When evolved in the heterogeneous (fluctuating) environment, smaller (FS) populations paid significant costs of adaptation across three distinct environmental pairs under consideration, but the larger (FL) populations avoided costs altogether. As described above, FS suffered significantly from T-G trade-offs while FL bypassed them. Interestingly, despite facing both T and G as the sole sources of carbon for equal number of generations (~120), the T-G trade-off manifested itself in FS as adaptation to T and maladaptation to G. To explain this asymmetry of fitness changes across T and G, we note that despite evolving in homogeneous G for ~480 generations, GS could not adapt significantly to this environment. Contrastingly, TS increased their fitness in T by > 1.5-fold within the same period (Fig. 3). This shows that the size of our small-population regimens was sufficient to adapt significantly to T but not to G. Put differently, the scope of adaptation in T was much greater than that in G (Chavhan *et al*. 2020). This can explain why FS adapted to T but not to G. Analogous to TS, such adaptation of FS to T also led to significant maladaptation in the other three environments.

In contrast to the small populations, the large populations in our study had sufficient supply of mutations to adapt to G within ~480 generations (Fig. 3). Curiously, we also found that FS could adapt significantly to G despite encountering this particular environment intermittently for a total period of ~120 generations (Fig. 2). This observation was also supported by the genome-wide analysis of the evolutionary changes in this regimen, which revealed substantial enrichment of putative G-associated beneficial mutations (Fig. 4).

An important alternative explanation for cost avoidance in heterogeneous environments involves the divergence of the population in question into multiple subpopulations, each one specialized on a different environmental component (Kassen 2002, 2014). However, our genomic data suggest that this explanation of cost avoidance in unlikely in our study. Specifically, in one of the sequenced FL populations (FL(a)), multiple mutations went to fixation, one of which was in a locus known to be associated with galactose uptake/metabolism and another with that of thymidine (Fig. 4). Hence, the individuals in FL(a) simultaneously carried both putative G and putative T adaptations. In the second sequenced FL population (FL(b)), a putative G-associated mutation went to fixation and three putative T-associated mutations rose to the frequencies of 60.7%, 41.4%, and 32.3% respectively (Fig. 4, Table S9). Thus, the probability that an individual in FL(b) carried at least one of the three putative T mutations was 84.41%. Hence, the probability that a given individual in FL(b) simultaneously carried a G- and T-associated mutation was 84.41%. Although the third sequenced FL population (FL(d)) did not show any fixation events, it enriched a putatively G-associated mutation at 57.70% and multiple putatively T-associated mutations at 68.5%, 63.0%, and 30.9%, respectively (Fig. 4, Table S9). Hence, the probability that an individual in this population carried at least one T-associated mutation was 91.95%. Moreover, the probability of an FL(d) individual simultaneously carrying both G- and T-associated mutations is 53.05%. Thus, the high likelihoods of simultaneously showing G- and T-associated mutations FL suggests that it is unlikely that this regimen avoided costs by divergent specialization on individual carbon sources within populations.

Overall, these results demonstrate that the phenomenon of cost avoidance in heterogeneous (fluctuating) environments requires the supply of variation to be large enough to make use of multiple dynamic selection pressures.

### Implications

Our observations offer a novel explanation for an important conundrum in evolutionary ecology, namely the rarity of detectable fitness costs in empirical studies. Specifically, we demonstrate a previously unreported interaction of population size and environmental heterogeneity that determines the evolutionary appearance (or avoidance) of fitness costs. These results can potentially explain how evolving populations can escape fitness costs despite substantial antagonistic pleiotropy across environmental states. Our study shows that the simultaneity of two conditions, namely large population size and heterogeneous environment, can avoid all the fitness costs that potentially evolve when these conditions are not present together. Finally, to our knowledge, this is the first experimental study to demonstrate that multiple mutations can fix rapidly (within ~480 generations) in asexual populations evolving in highly dynamic heterogeneous environments, a possibility raised recently (Cvijović *et al*. 2015), but discounted by older studies (Whitlock 1996; Kassen 2002). Remarkably, this phenomenon was observed in both FL and FS populations. This shows that such rapid fixation of multiple mutations in heterogeneous environments can happen in the face of both lenient and harsh population bottlenecks.

The environments of most natural populations of asexual microbes are known to be heterogeneous (Green & Bohannan 2006; Muscarella *et al*. 2019). Moreover, such natural asexual populations are also known to have extremely large sizes (Torsvik *et al*. 2002; Tenaillon *et al*. 2010). Our results suggest that if the asexual population under consideration has a history of evolving in heterogeneous environments in large numbers, it is expected to have reached its current state after having avoided fitness costs during its past evolution. Therefore, if a sample from such a population is now employed to analyse fitness correlations in a single-generation study, such correlations may not be negative, and costs may not be found.

Contrastingly, several laboratory evolution studies using unchanging (homogeneous) environments and large population sizes (> 10^6^ in terms of harmonic mean population size) have successfully detected fitness costs (Kassen & Bell 1998; Cooper & Lenski 2000; Cooper *et al*. 2001; Nilsson *et al*. 2004; Hall & Colegrave 2008; Presloid *et al*. 2008; Philippe *et al*. 2009; Vasilakis *et al*. 2009; Bedhomme *et al*. 2012; Ensminger *et al*. 2012; Kubinak & Potts 2013; Leiby & Marx 2014). This agrees with the interplay of population size and environmental heterogeneity revealed by our results, which predicts such a combination of constant environment and large populations to lead to significant costs.

Thus, apart from explaining why costs may not be detected in single-generation studies with natural isolates, our observations also explain why costs can still be detected if the artificially controlled laboratory conditions remain constant over a few hundred generations in an evolution experiment.

Although the environments used in our experimental setup were nutritionally challenging minimal media, the explanation of our observations applies to the general notion of fitness costs across multiple environments in asexual microbial populations. In particular, our results can have important implications for understanding the rampant evolution and spread of antibiotic resistance, which has direct practical values. Mutations that confer resistance to antibiotics have been routinely shown to bear fitness costs in drug-free conditions (Andersson & Hughes 2010; Vogwill & MacLean 2015). Interestingly, resistant microbes mostly evolve in a heterogeneous environment that fluctuates randomly across antibiotic-laden and antibiotic-free conditions (Baquero *et al*. 1998). Our results predict that small populations evolving in heterogeneous environments suffer heavy fitness costs while large populations are likely to avoid them altogether (Fig. 2). Thus, even if most antibiotic resistance mutations carry a cost in drug-free conditions, large microbial population sizes stemming from lack of sanitary conditions and proper medical waste-disposal (Cantón *et al*. 2013) could themselves lead to vigorous spread of cost-free resistance.

## Methods

### Experimental evolution

We derived ten different evolutionary regimens from a single colony of *E. coli* MG1655 by culturing populations at two different sizes in five different environments as described above (see Supplementary Methods (SM.1) for more details regarding the ancestral strain and media compositions). Using the standard batch culture technique, we let all the 60 populations propagate as continuously shaken cultures (150 rpm) in 96 well plates maintained at 37° C. In all the 60 populations, the culture volume was fixed at 300 μl. Whereas the large (L) populations experienced a lenient periodic bottleneck (1:10, the small (S) populations faced a relatively harsher periodic bottleneck (1:10^4^ dilution). We ensured that populations of different sizes did not remain in the stationary phase for significantly different time-periods by bottlenecking the L populations every 12 hrs (~3.3 generations), and the smaller ones every 48 hrs (~13.3 generations). The selection protocol pertaining to the T and G populations has been reported in a previous study (Chavhan *et al*. 2020).

### Fitness quantification

We conducted fitness measurements for all the 60 independently evolving populations in all four carbon sources (T, G, M, and S) at the end of the evolution experiment (~480 generations). To this end, we revived the cryo-stocks belonging to each of the 60 experimental populations in a common nutrient limited environment that was not encountered by any population during the ~480 generations of our experiment (glucose based M9 minimal medium) and allowed them to grow for 24 hours. Using a well-plate reader (Synergy HT, BIOTEK^®^ Winooski, VT, USA), we then performed automated growth measurements on each of the 60 revived populations in all four different minimal media, each based on one of T, G, M, or S. Ensuring that the physical conditions during the fitness measurements were the same as the culture conditions (96 well plates shaken at 150 rpm and ambient temperature maintained at 37° C), we obtained growth readings every 20 minutes for 24 hours. We used optical density (OD) at 600 nm as the measure of population density.

Since the total number of growth curves was much larger than number of wells in the assay plate, we used a randomized complete block design (RCBD) for growth measurements(Milliken & Johnson 2009). Specifically, we assayed one replicate population of each of the ten different evolutionary lines in all four environments on a given day. Since there were six replicates for each evolutionary line, we conducted growth measurements over six different days. We used the maximum growth rate (R) as the measure of fitness. We computed R as the maximum slope of the growth curve over a dynamic window of ten OD (600 nm) readings (Leiby & Marx 2014; Karve *et al*. 2015; Chavhan *et al*. 2019a, b). As described in the Results section, for each of the four sole carbon sources (G, M, S, and T), we used single sample t-tests to compare the fitness of each of the ten evolutionary regimens to that of the ancestor. Subsequently, we corrected for family-wise error rates using the Holm-Šidàk procedure.

As described in the Supplementary Methods, we also investigated the changes in the geometric mean fitness across G, M, S, and T for all the ten evolutionary regimens (see SM.2 for details).

### Whole genome whole population sequencing

For both the ancestor and the six randomly chosen evolved populations (three each from FL and FS), pellets obtained from overnight grown cultures were sent for sequencing to an external service provider. For each sample, the genomic DNA was isolated using c-TAB and phenol-chloroform extraction. This procedure was followed by RNAase A treatment. The quality and quantity of the isolated DNA samples was verified using a NanoDrop™ spectrophotometer (Thermo Fisher Scientific Inc., MA, USA). The isolated DNA samples were initially subjected to a further check by targeting the bacterial 16s gene using Sanger sequencing. After these checks, 2 x 150 NextSeq500 Shotgun Libraries were prepared from each sample using an Illumina TruSeq^®^ Nano DNA Library Prep Kit (Illumina Inc, CA, USA). The quality of each library was checked using the Agilent 4200 Tape Station (Agilent Technologies, CA, USA). The libraries were then loaded onto NextSeq500 (Illumina Inc, CA, USA) for cluster generation and paired-end sequencing. Trimmomatic (v0.38) was used to remove adapter sequences, ambiguous reads (with unknown nucleotides > 5%) and low-quality sequences (reads with > 10% quality threshold < 20 phred score). After trimming, a minimum length of 100nt was applied. The mean coverage across the sequenced populations was ~100-fold at a quality score of 20.

We subjected these trimmed high quality sequences to the BRESEQ pipeline (Deatherage & Barrick 2014) (v0.33.2) to identify mutations enriched during our evolution experiment. We initially compared the ancestral sequence to the reference *E. coli* MG1655 genome to identify differences relative to the latter expected to be found in all the six evolved populations. Next, we adjusted for these differences by using the ancestral sequences as the reference for identifying mutational frequencies in each of the six descendant populations using the ‘polymorphic’ mode in BRESEQ. To avoid false positives and to restrict our analysis to mutations that must have been instrumental in shaping the average fitness of the population, we ignored mutations with frequencies < 10%.

## Acknowledgements

We thank S. Selveshwari for help with NGS analysis and Milind Watve and M.S. Madhusudhan for their valuable inputs. YDC was supported by a Senior Research Fellowship initially sponsored by IISER Pune and then by Council for Scientific and Industrial Research (CSIR), Govt. of India. YDC also acknowledges Department of Biotechnology, Govt. of India for postdoctoral funding for this work. SM was supported by an INSPIRE undergraduate fellowship, sponsored by Department of Science and Technology (DST), Govt. of India. This project was supported by an external grant (BT/PR22328/BRB/10/1569/2016) from Department of Biotechnology, Govt. of India, and internal funding from IISER Pune.

## Data archiving

All the data relevant to this study will be uploaded on the Dryad digital repository upon acceptance.

## Supplementary Information

**Table S1.**
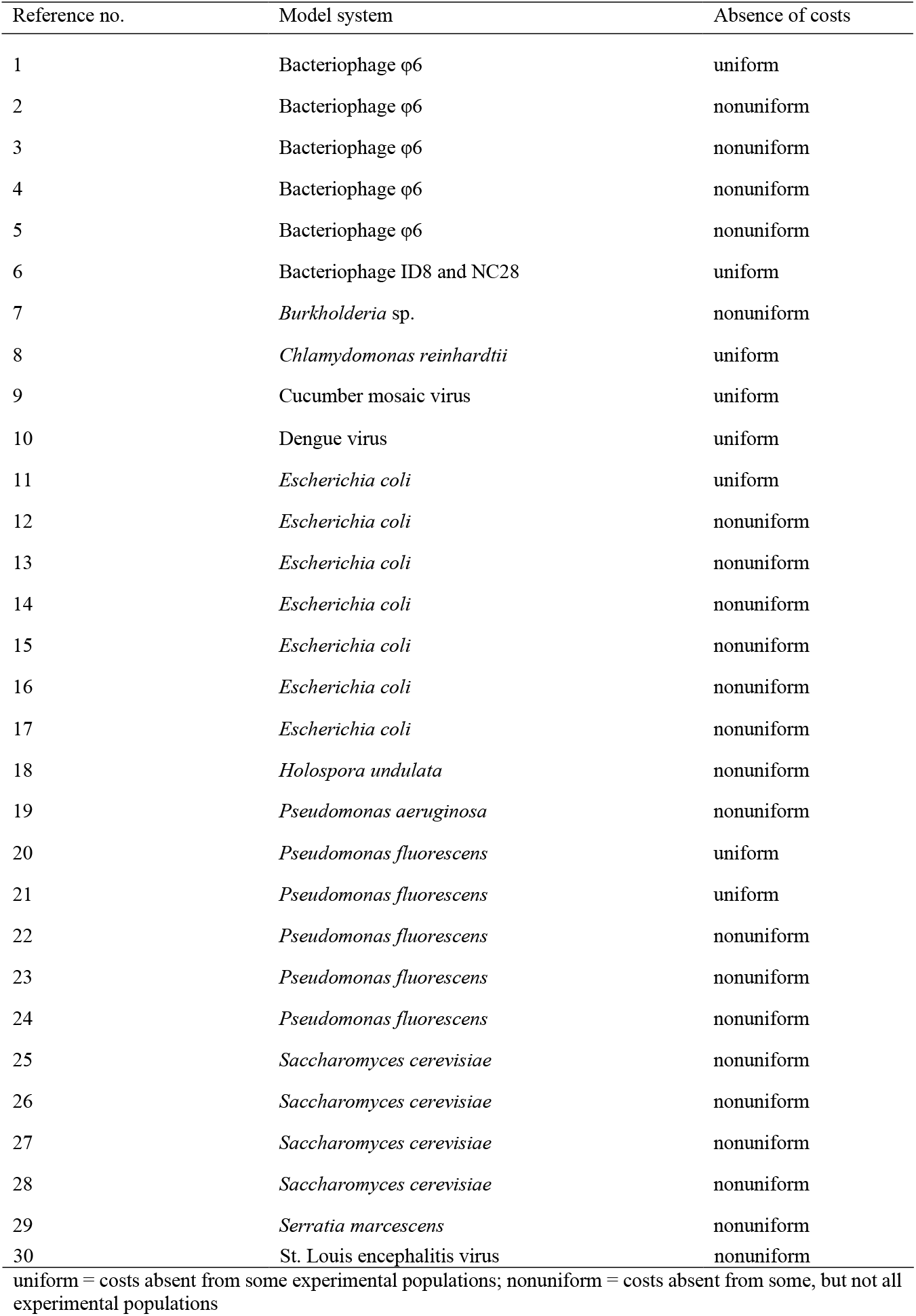
The absence of costs in experimental evolution studies with asexual microbes

### Supplementary text

#### ST.1. Details of the studies shown in Fig. 1a

Fig. 1a incorporates those bacterial experimental evolution studies on fitness costs in heterogeneous environments for which estimates of harmonic mean population size could be obtained. Studies conducted with viruses, and eukaryotes are not included here.

Key in the legend of Fig. 1a:

A: Ref. 29*; B: Ref. 24; C: Ref. 19; D: Ref. 23; E: Ref. 15; F: Ref. 11; G: Ref. 22; H: Ref. 20; I: Ref. 17^‡^

*The population size reported for Study A (Ref. 29) has been calculated indirectly using the stationary phase densities reported for a different bacterial species in the selection medium in question and is likely an overestimate.

‡The data for population size have been provided by the authors of Study I (Ref. 17).

### Supplementary Methods

#### SM.1. Details of the ancestral strain and nutrient media

##### Ancestral strain

*Escherichia coli* MG1655 lacY::kan. The ancestral strain was resistant to kanamycin.

##### Nutrient media

There was one heterogeneous and four homogeneous environments in our evolution experiment. Each homogeneous environment comprised of an M9-based minimal medium, 1 litre of which contained the following:

- 12.8 g Na_2_HPO_4_.7H_2_O
- 3.0 g KH_2_PO_4_
- 0.5 g NaCl
- 1.0 g NH_4_Cl
- 240.6 mg MgSO_4_
- 11.1 mg CaCl_2_
- 4g of the pre-decided sole carbon source
- 50 mg Kanamycin sulphate

The four homogeneous environments differed in terms of the identity of the pre-decided sole carbon source. The following four carbon sources were used in our experiment:

- Thymidine
- Galactose
- Maltose
- Sorbitol

The heterogeneous environment fluctuated randomly between the above four carbon sources every 13.3 generations.

#### SM.2. Analysis of differences in geometric mean fitness in our experimental regimens

We computed the geometric mean fitness across each of the four carbon sources (G, M, S, and T) for all the ten evolutionary regimens in our experiment (comprising sixty independently evolving populations in total).

We used a mixed model ANOVA to compare the geometric mean fitness across the populations evolved in the heterogeneous environment (FL and FS). In this analysis, we considered the population size (two levels: large (L) and small (S)) as the fixed factor and the day of assay as the random factor, with each day corresponding to one biological replicate in our randomized complete block design (RCBD (see the Main text for details)). We also determined the effect size of the difference between FL and FS using partial η^2^, interpreting the latter as showing small, medium, or large effect for Partial η^2^ < 0.06, 0.06 < Partial η^2^ < 0.14, 0.14 < Partial η^2^ respectively^31^.

We further tested if the treatment regimens evolved in the heterogeneous environment (FL / FS) had evolved significantly different geometric mean fitness (over T, G, M, and S) as compared to the control regimens evolved in homogeneous environments. To this end, we conducted two mixed-model ANOVAs with evolutionary regimen (nine levels) as the fixed factor and day of assay (six levels) as the random factor. In the first ANOVA (Table S6), the nine levels in the evolutionary regimen (fixed factor) consisted of the eight homogeneous environments regimens and FL, while in the second ANOVA (Table S7), the fixed factor consisted of the same eight homogeneous environments and FS. For both ANOVAs, we used the Dunnett’s procedure) to assess the pairwise differences of FL or FS with the eight homogeneous environment regimens.

In another (more conservative) analysis of the differences in GM fitness across regimens, we used a mixed model ANOVA with evolutionary regimen (ten levels: FL, FS and eight homogeneous environment regimens) as the fixed factor and day of assay (six levels) as the random factor. Subsequently, we compared all possible pairwise differences between the ten evolutionary regimens using Tukey’s HSD (Table S8).

### Supplementary Results

**Table S2.** Analysis of adaptation and maladaptation events in all ten evolutionary regimens using single-sample t-tests (N = 6) with reference to the ancestral fitness in each of the four carbon sources* (scaled to 1).

| Selection environment | Population type | Assay environment | $P$ value | Corrected $P$ value | Inference |
| --- | --- | --- | --- | --- | --- |
| Heterogeneous | FL | T | $1.58 \times 10^{-5}$ | $6.33 \times 10^{-5}$ | Adaptation |
| Heterogeneous | FL | G | $7.025 \times 10^{-3}$ | 0.021 | Adaptation |
| Heterogeneous | FL | M | 0.564 | - | No change |
| Heterogeneous | FL | S | 0.612 | - | No change |
| Heterogeneous | FS | T | 0.049 | 0.049 | Adaptation |
| Heterogeneous | FS | G | 0.025 | 0.051 | Maladaptation |
| Heterogeneous | FS | M | $2.5 \times 10^{-4}$ | 0.001 | Maladaptation |
| Heterogeneous | FS | S | 0.009 | 0.02635 | Maladaptation |
| Homogeneous T | TL | T | $8.1 \times 10^{-7}$ | $3.24 \times 10^{-6}$ | Adaptation |
| Homogeneous T | TL | G | $8.94 \times 10^{-5}$ | $2.68 \times 10^{-4}$ | Maladaptation |
| Homogeneous T | TL | M | $5.53 \times 10^{-4}$ | $5.53 \times 10^{-4}$ | Maladaptation |
| Homogeneous T | TL | S | $1.13 \times 10^{-4}$ | $2.27 \times 10^{-4}$ | Maladaptation |
| Homogeneous T | TS | T | $1.83 \times 10^{-4}$ | $7.33 \times 10^{-4}$ | Adaptation |
| Homogeneous T | TS | G | $6.839 \times 10^{-3}$ | $6.839 \times 10^{-3}$ | Maladaptation |
| Homogeneous T | TS | M | $4.58 \times 10^{-4}$ | $1.372 \times 10^{-3}$ | Maladaptation |
| Homogeneous T | TS | S | $6.19 \times 10^{-4}$ | $1.238 \times 10^{-3}$ | Maladaptation |
| Homogeneous G | GL | T | 0.003 | 0.013 | Maladaptation |
| Homogeneous G | GL | G | 0.016 | 0.049 | Adaptation |
| Homogeneous G | GL | M | 0.633 | 0.633 | No change |
| Homogeneous G | GL | S | 0.973 | 0.973 | No change |
| Homogeneous G | GS | T | 0.122 | 0.122 | No change |
| Homogeneous G | GS | G | 0.617 | 0.617 | No change |
| Homogeneous G | GS | M | 0.483 | 0.483 | No change |
| Homogeneous G | GS | S | 0.016 | 0.061 | No change |

**Table S2 continued**
| Selection environment | Population type | Assay environment | *P* value | Corrected *P* value | Inference |
| --- | --- | --- | --- | --- | --- |
| Homogeneous M | ML | T | 0.252 | - | No change |
| Homogeneous M | ML | G | 0.036 | 0.134661 | No change |
| Homogeneous M | ML | M | 0.090 | - | No change |
| Homogeneous M | ML | S | 0.762 | - | No change |
| Homogeneous M | MS | T | 0.134 | - | No change |
| Homogeneous M | MS | G | 0.164 | - | No change |
| Homogeneous M | MS | M | 0.066 | - | No change |
| Homogeneous M | MS | S | 0.069 | - | No change |
| Homogeneous S | SL | T | $3.14 \times 10^{-4}$ | $1.257 \times 10^{-3}$ | Maladaptation |
| Homogeneous S | SL | G | $7.59 \times 10^{-3}$ | 0.023 | Maladaptation |
| Homogeneous S | SL | M | 0.088 | - | No change |
| Homogeneous S | SL | S | 0.690 | - | No change |
| Homogeneous S | SS | T | $5.5 \times 10^{-3}$ | 0.011 | Maladaptation |
| Homogeneous S | SS | G | $6.4 \times 10^{-5}$ | $2.56 \times 10^{-4}$ | Maladaptation |
| Homogeneous S | SS | M | 0.001 | 0.003 | Maladaptation |
| Homogeneous S | SS | S | 0.366 | - | No change |

\*The data pertaining to evolution in T and G have been reported in a previous study<sup>32</sup>.

**Table S3.** The evolutionary emergence of costs of adaptation in populations evolved in the heterogeneous environment

| Case(s) which showed costs of adaptation |  | Case(s) which showed simultaneous adaptation |  |
| --- | --- | --- | --- |
| Population size | Environmental pair(s) | Population size | Environmental pair(s) |
| Large | None | Large | T (adaptation) – G (adaptation) |
| Small | T (adaptation) - G (maladaptation) | Small | None |
|  | T (adaptation) - S (maladaptation) |  |  |
|  | T (adaptation) - M (maladaptation) |  |  |

**Table S4.** The evolutionary emergence of costs of adaptation in populations evolved in homogeneous environments

| Case(s) which showed costs of adaptation |  |  | Case(s) which showed simultaneous adaptation |
| --- | --- | --- | --- |
| Selection environment | Population size | Environmental pair(s) |  |
| T | Large | T (adaptation) - G (maladaptation)<sup>†</sup> | None |
|  |  | T (adaptation) - S (maladaptation)<sup>†</sup> |  |
|  |  | T (adaptation) - M (maladaptation)<sup>†</sup> |  |
| T | Small | T (adaptation) - G (maladaptation) |  |
|  |  | T (adaptation) - S (maladaptation) |  |
|  |  | T (adaptation) - M (maladaptation) |  |
| G | Large | G (adaptation) - T (maladaptation) |  |
| G | Small | Not applicable<br>(no adaptation to the selection environment) |  |
| S | Large |  |  |
| S | Small |  |  |
| M | Large |  |  |
| M | Small |  |  |

<sup>†</sup>Across the T - M, T - G, and T - S pairs, the costs suffered by the large (TL) populations were greater than those suffered by the small (TS) populations<sup>32</sup>.

**Fig. S1.**
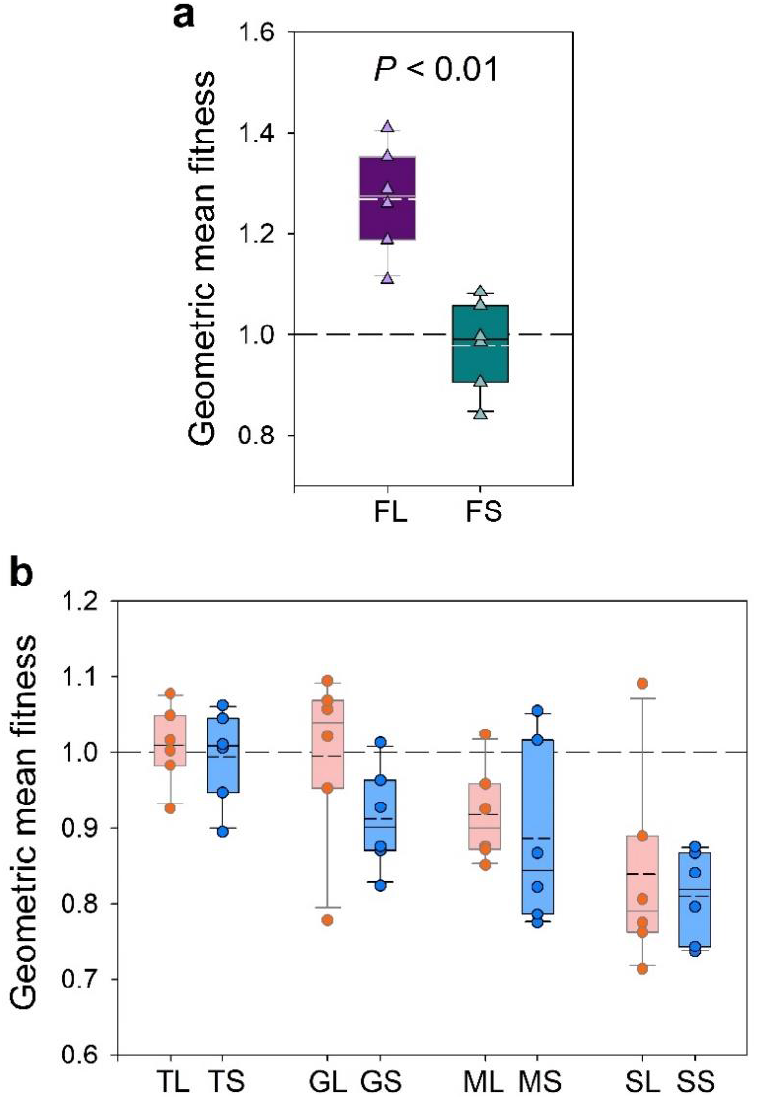
Changes in geometric mean fitness of our experimental populations across T, G, M, and S. The solid lines in the box plots mark the 25^th^, 50^th^, and 75^th^ percentiles while the whiskers mark the 10^th^ and 90^th^ percentiles; the short-dashed lines within the box plots represent means (N = 6). The long-dashed line outside the box plots represent the ancestral level of the ordinate. **(a)** Geometric mean fitness of populations evolved in the heterogeneous environment. FL > FS (P < 0.01). **(b)** Geometric mean fitness of populations evolved in homogeneous environments. See Tables S5 and S6 for details.

We found that FL populations had significantly higher geometric mean fitness than FS (Fig. 5a; Table S4; mixed-model ANOVA: F1,5 = 18.002; *P* = 0.008; partial η^2^ = 0.783 (large effect)).

Thus, the large (FL) populations adapted better than the small (FS) populations to their common heterogeneous environment. This result is expected from the absence of any fitness costs in FL and the presence of such costs across the maximum possible number of environmental pairs under consideration in FS. We further found that FL could significantly enhance their geometric mean fitness with respect to the common ancestor, but FS failed to do so (Fig. S1a; Table S5). Curiously, despite showing significant fitness changes in T (adaptation) and G (maladaptation), FS did not have significantly different geometric mean fitness as compared to the common ancestor (Fig. S1a; Table S5). Adaptation to homogeneous environments is not expected to entail increased geometric mean fitness over multiple (unencountered) environments. Indeed, we found that the geometric mean fitness over the four carbon sources did not increase significantly as compared to the ancestral level in any of the homogeneous environment regimens, regardless of the population size (Fig. S1b; Table S5).

We also found that FL had a much larger geometric mean fitness than all the homogeneous environment regimens (Table S6). However, FS did not have significantly different geometric mean fitness as compared to a vast majority (seven out of the eight) of homogeneous environment regimens (Table S7). A similar pattern was revealed by a more conservative post hoc analysis using Tukey’s HSD (Table S8).

Both the above analyses (using Dunnett’s or Tukey’s post-hoc tests) sought to answer the same question: whether the FL / FS regimens significantly differed in their GM fitness as compared to the homogeneous environment regimens. Comparing Tables S6–S7 with S8, we find that the pair-wise differences that turn up as statistically significant are identical between the two analyses (except one case: FS and SL show up as significantly different in Dunnett’s test but not in Tukey’s HSD). This is not surprising, as the analysis with two Dunnett’s procedures comprises of (and therefore corrects for) only 18 pair-wise tests, while the corresponding analysis with Tukey’s HSD corrects for 81 pair-wise tests (of which only 18 are relevant for our purpose). Therefore, the second analysis has a lot less power than the first one. The fact that the results remain virtually identical across both cases highlights the robustness of the same. It should be noted here that our interpretation of the difference between FL and FS remains agnostic to the choice of analysis.

Taken together, FL adapted significantly to the heterogeneous (fluctuating) environment, but FS failed to do so. Importantly, the preparedness of FS to face the environmental fluctuations across G, M, S and T was similar to most homogeneous environment regimens.

**Table S5.** Summary of single-sample t-tests (N = 6) of differences in the geometric mean fitness (calculated over G, M, S, and T) of the ten evolutionary regimens with the corresponding ancestral value (= 1)

| Selection environment | Population type | $P$ value | Inference |
| --- | --- | --- | --- |
| Heterogeneous | FL | 0.002 | GM enhanced |
| Heterogeneous | FS | 0.584 | No change |
| Homogeneous | TL | 0.703 | No change |
| Homogeneous | TS | 0.826 | No change |
| Homogeneous | GL | 0.922 | No change |
| Homogeneous | GS | 0.026 | GM reduced |
| Homogeneous | ML | 0.027 | GM reduced |
| Homogeneous | MS | 0.069 | No change |
| Homogeneous | SL | 0.034 | GM reduced |
| Homogeneous | SS | $5.96 \times 10^{-4}$ | GM reduced |

**Table S6.** Summary of Dunnett post-hoc tests (N = 6) with respect to FL done after analysing the geometric mean fitness differences across nine evolutionary regimens (FL and eight homogeneous environment regimens) using a mixed model ANOVA, which revealed a significant main effect of the identity of the evolutionary regimen: F8,40 = 16.284, *P* = 2.172 × 10^−10^

| Population type | $P$ value (Dunnett (reference: FL)) |
| --- | --- |
| GL | 0.000016 |
| GS | 0.000009 |
| TL | 0.000028 |
| TS | 0.000016 |
| ML | 0.000009 |
| MS | 0.000009 |
| SL | 0.000009 |
| SS | 0.000009 |

**Table S7.** Summary of Dunnett post-hoc tests (N = 6) with respect to FS done after analysing the geometric mean fitness differences across nine evolutionary regimens (FS and eight homogeneous environment regimens) using a mixed model ANOVA, which revealed a significant main effect of the identity of the evolutionary regimen: F8,40 = 5.094, *P* = 2.074 × 10^−4^

| Population type | *P* value (Dunnett (reference: FS)) |
| --- | --- |
| GL | 0.999771 |
| GS | 0.577267 |
| TL | 0.984290 |
| TS | 0.999855 |
| ML | 0.666052 |
| MS | 0.243947 |
| SL | 0.024777 |
| SS | 0.004229 |

**Table S8.** Summary of Tukey post-hoc tests (N = 6) done after analysing the geometric mean fitness differences across all the ten evolutionary regimens using a mixed model ANOVA, which revealed a significant main effect of the identity of the evolutionary regimen: F9,45 = 14.566, *P* = 1.129 × 10^−10^. Tukey *P* values for pairwise differences with only FL and FS are shown below:

|  | GL | GS | TL | TS | ML | MS | SL | SS | FL | FS |
| --- | --- | --- | --- | --- | --- | --- | --- | --- | --- | --- |
| FL | 0.000177 | 0.000156 | 0.000216 | 0.000175 | 0.000156 | 0.000156 | 0.000156 | 0.000156 | - | 0.000161 |
| FS | 0.999998 | 0.921270 | 0.999688 | 0.999999 | 0.952902 | 0.647098 | 0.126786 | 0.027274 | 0.000161 | - |

**Table S9.**
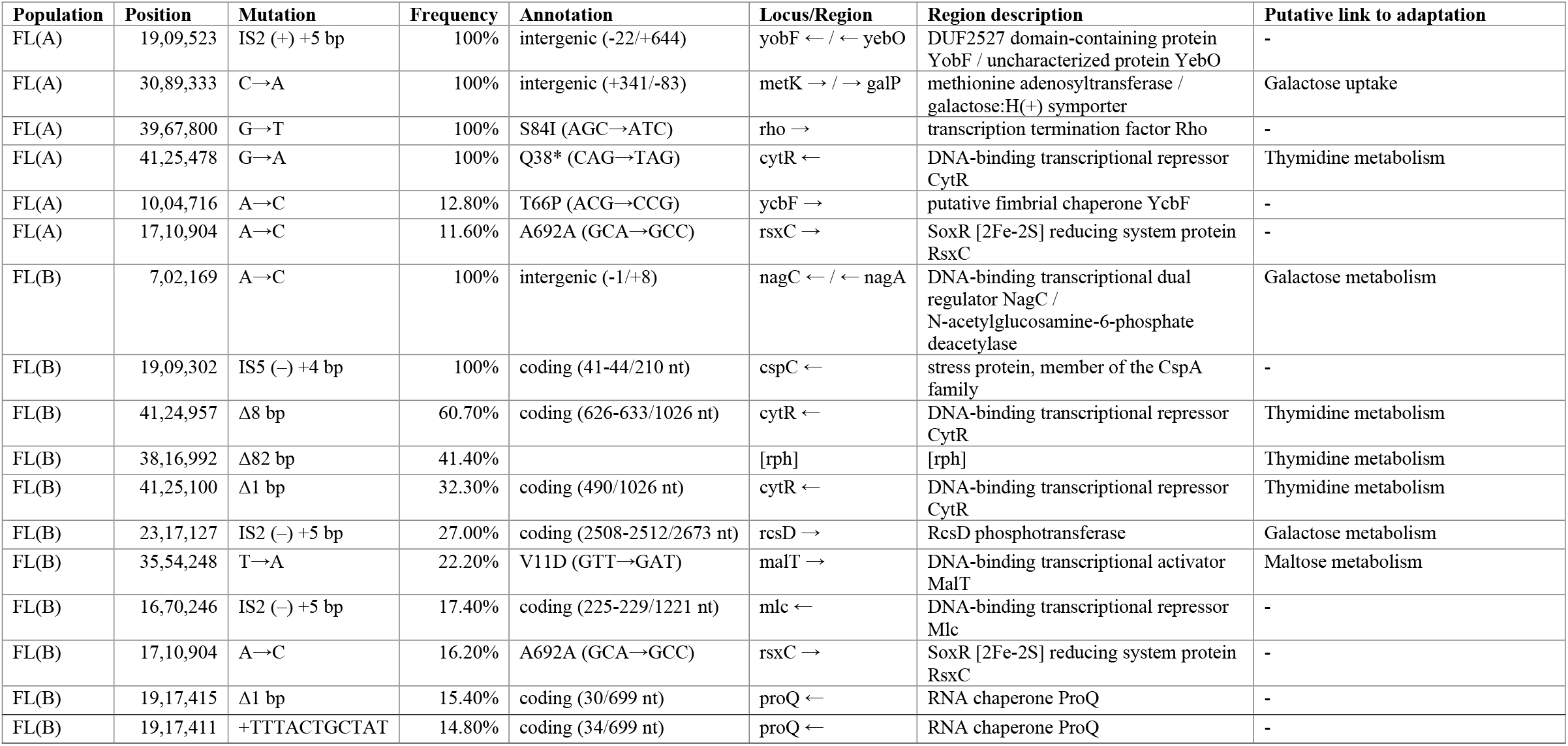
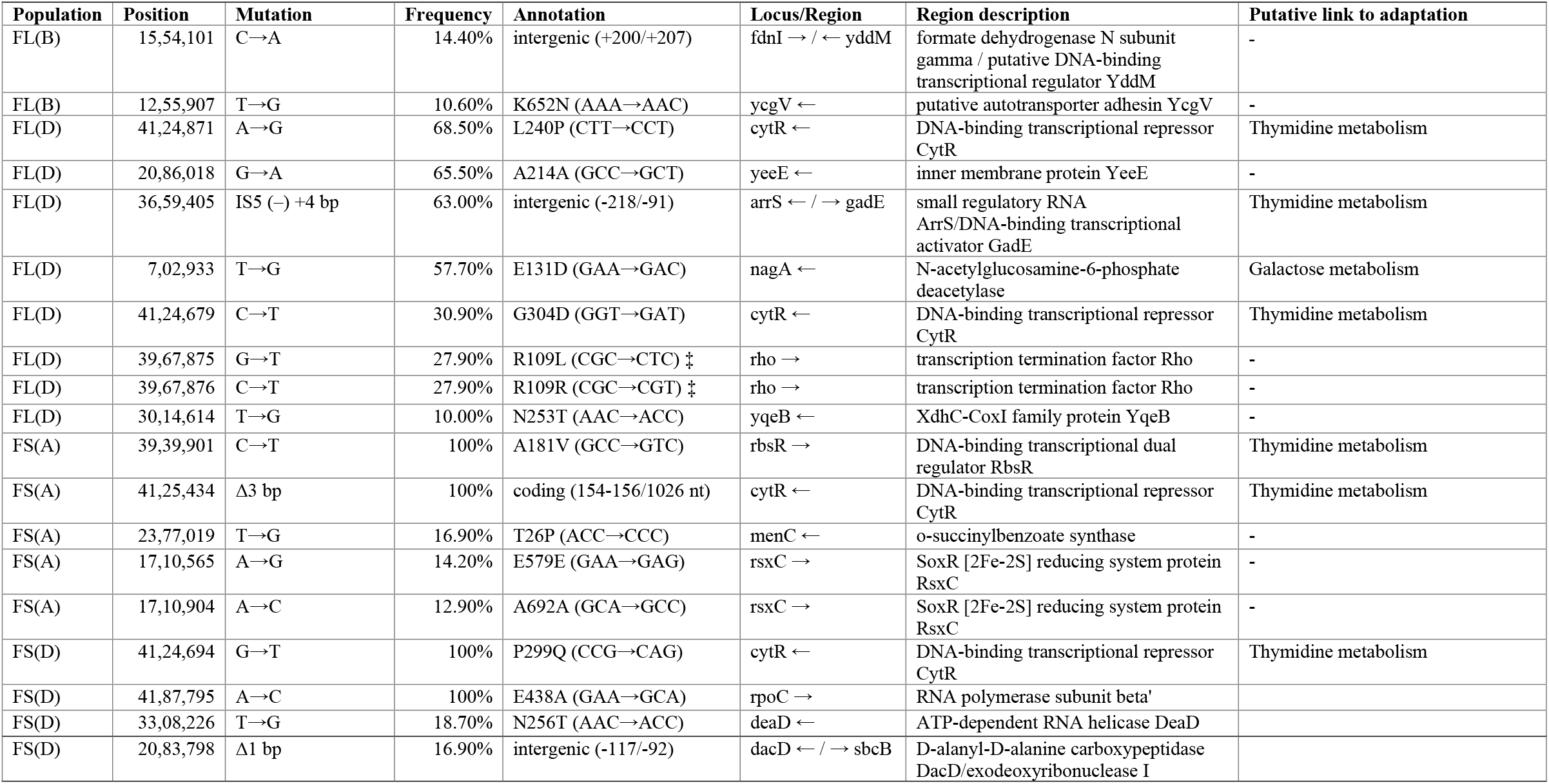
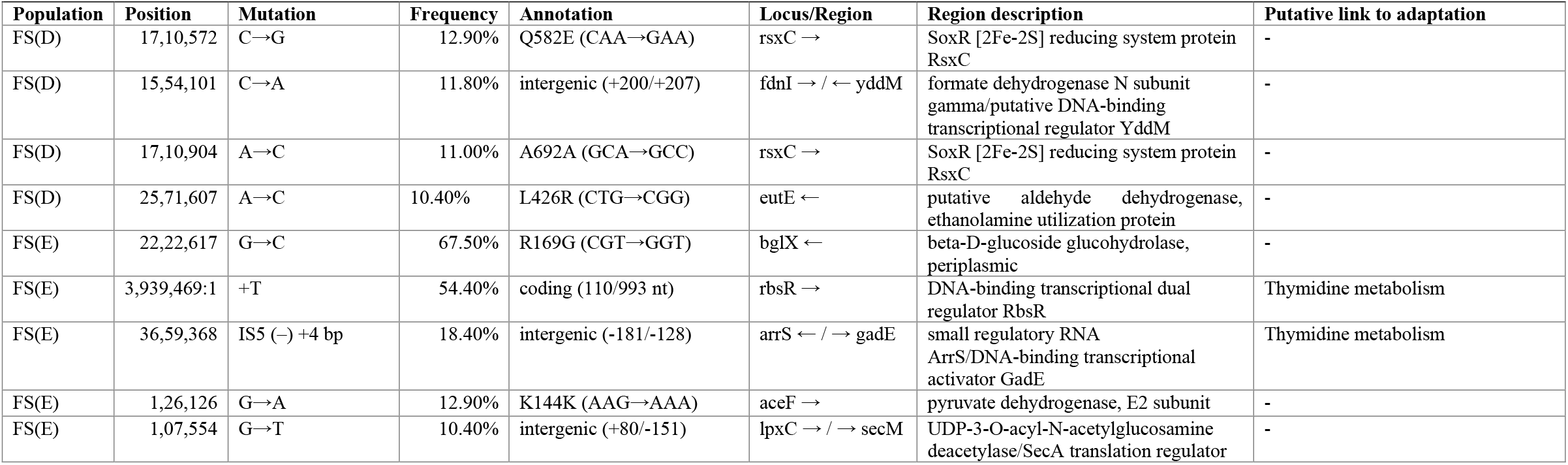
Details of mutations observed at frequencies ≥ 10% after ~480 generations of evolution

